# Decellularized Small Intestine for Full Thickness Burn Wound Treatment

**DOI:** 10.64898/2026.02.10.704815

**Authors:** Inês V. Silva, Ilda Rodrigues, Clara Sousa, Raquel Costa, Lorenzo Moroni, Ana L. Oliveira

**Affiliations:** Universidade Católica Portuguesa, CBQF - Centro de Biotecnologia e Química Fina – Laboratório Associado, Escola Superior de Biotecnologia, Porto, Portugal; Maastricht University, MERLN Institute for Technology-Inspired Regenerative Medicine, Complex Tissue Regeneration Department, 6229 ER, Maastricht, the Netherland; Biochemistry Unit, Department of Biomedicine, Faculty of Medicine, University of Porto, Porto, Portugal; i3S-instituto de inovação e investigação em saúde, University of Porto, Portugal

**Keywords:** decellularization, extracellular matrix, small intestine, wound regeneration, full thickness burn

## Abstract

Treating extensive full-thickness burn wounds remains difficult in clinical practice because available donor skin is often limited, the risk of infection is high, and many standard dressings do not perform well when defects are large or structurally complex. These limitations have shifted attention to decellularized extracellular matrix (dECM) scaffolds, which can provide physical coverage while preserving biochemical cues that may support tissue repair. Based on this rationale, we designed a decellularization method that improves reagent penetration to produce a full-thickness porcine decellularized small intestine (dSI) scaffold for use in burn wound coverage. The protocol removed most cellular material while leaving low levels of detergent residue, and it maintained the native three-layer structure of the intestinal wall. Most key ECM components, such as collagen and glycosaminoglycans, were also retained. In this study, the dSI showed several properties relevant to burn care, capacity to absorb large amounts of fluid, water vapor transmission rates similar to those reported for skin, and resisted microbial penetration *in vitro*. From a mechanical standpoint, the scaffold retained anisotropic behaviour and remained stable under cyclic loading. This pattern indicates that it could withstand repeated deformation instead of acting like a fragile membrane. Degradation tests under enzymatic and oxidative conditions indicate that the material breaks down in a controlled way over a period that appears consistent with typical wound-healing timelines. *In vitro* assays indicated that the scaffold was cytocompatible, as human dermal fibroblasts and keratinocytes both attached to its surface and continued to proliferate. Cell responses differed depending on surface orientation, suggesting that preserved intestinal layers may shape cell behaviour in ways that are often missing in thinner or more uniform matrices. Overall, full-thickness dSI appears to act as a biologically active scaffold and shows mechanical properties that exceed those of many currently used burn dressings.

## 1. Introduction

Burn injuries are a major global health challenge, affecting millions of people worldwide. It is estimated that there are approximately 11 million burn injuries and 180,000 fatalities annually, making it a significant public health concern (1). The morbidity of burn injuries extends far beyond the initial physical trauma, as the resulting complications can be severe and long-lasting (1). Beyond the immediate damage to the skin and underlying tissues, burn injuries often lead to microorganism invasion, that can quickly turn into infection, which then can result in the onset of sepsis, a life-threatening condition (1). Moreover, the long-term effects of burn scarring can have a profound impact on a person’s quality of life. Burn scars can compromise joint mobility, limit functionality, hinder the ability to perform routine daily tasks, and cause permanent nerve damage which in turn can create chronic pain, hypersensitivity and loss of sensation. This can significantly impair an individual’s physical and emotional well-being, making it crucial to provide comprehensive care and support for burn survivors (2, 3). In the event of a burn, the standard treatment guide typically begins with cleaning the wound, controlling infection and stabilizing the patient. Moreover, severe burn wounds management may include the application of topical antimicrobial agents, surgical debridement, followed by skin grafting or the use of advanced wound dressings (4). The ideal burn wound dressing typically should maintain a moist environment while effectively managing exudate, act as a barrier against microbial invasion, permit gas exchange, and ensure it is non-toxic, biocompatible, and cost-efficient (5, 6). Conventional dressings and autografts, while presenting some of the necessary properties, face significant limitations in the treatment and healing of burn wounds, necessitating the development of novel therapeutic strategies (1).

Commercial burn dressings span a wide range of materials, including bandages, foams, films, and hydrogels. In routine clinical practice, cotton gauze remains one of the most frequently used options, largely because it is inexpensive and readily available. The need for frequent dressing changes is a notable drawback. Repeated removal of gauze can disturb newly formed tissue and, in some cases, lead to partial wound reopening. This risk becomes especially problematic in deeper burn injuries, where anatomical and functional restoration is already difficult and often prolonged (7).

Autografts are still widely regarded as the clinical gold standard for burn treatment, largely because of their excellent biocompatibility, low risk of immune rejection, and reliable ability to achieve wound closure. Their effectiveness, however, becomes less certain in the context of extensive burns. When large surface areas are affected, the amount of healthy donor skin available is often insufficient to meet clinical demands (8). In practical terms, the body may simply not be able to supply enough viable tissue to cover all affected regions. (8). Complicating matters further, harvesting skin for autografting inevitably creates a secondary wound at the donor site. This additional injury carries its own risks, including pain, infection, and delayed healing, and can place a significant burden on already vulnerable patients. In some cases, donor sites develop hypertrophic scarring or pigmentation abnormalities, outcomes that may persist long after the primary burn has healed and contribute to both physical discomfort and psychological distress (8). Taken together, these limitations help explain why alternative strategies and adjunctive therapies continue to attract considerable attention in the field of burn care, to reduce morbidity while improving long-term outcomes.

Given these limitations, hydrogels have emerged as a potentially more suitable class of commercial dressings (7). Their excellent swelling ability and moisture retention appears to support cell migration and tissue regeneration. Beyond their passive physical properties, hydrogels can also be modified or combined with additional components to address specific clinical needs. For instance, antimicrobial agents are often incorporated to reduce bacterial burden at the wound site, an issue that remains a persistent concern in burn care (7). Hydrogels adaptability has made them increasingly attractive compared to more traditional dressings. However, they typically lack the complex biological cues present in native extracellular matrix (ECM), such as adhesion ligands, growth factors, and structural proteins, which are crucial for guiding cell migration, proliferation, and tissue regeneration (9).

Xenographic tissue, after adequate decellularization processing cope with the low immunogenicity requirements, representing a unique avenue for developing advanced wound dressings (10). Porcine small intestine submucosa is often used to generate scaffolds for wound treatment application due to its composition of fibroblast growth factors, transforming growth factor-beta, vascular endothelial growth factor, and structural and functional proteins (e.g. fibronectin, glycosaminoglycan and hyaluronic acid) (11). These components play pivotal roles in wound healing, regulating cell division, migration, and differentiation (12). Moreover, the three-dimensional structure of the decellularized extracellular matrix (dECM) offers a well-organized and bioactive surface that is rich in receptors facilitating cell attachment. This structural organization closely mimics the native cellular environment, providing essential mechanical stability that supports cell proliferation and migration (12). This combination of physical support and biochemical stimuli makes this dECM an promising scaffold for wound healing to restore impaired dynamic reciprocity mechanism (12).

To fully preserve these important bioactive molecules while ensuring its cost-effectiveness is an essential task, that can only be achieved by adequately designing tissue specific decellularization processes. Moreover, the three layer system of the small intestine hasn’t been extensively explore has a more relevant scaffold to produce a skin mimetic for full thickness burn wounds against commercial single layer dECM (e.g. amniotic membrane and small intestine submucosa) that tend to be mechanically weaker, degraded unpredictably which in turn compromises cell attachment and healing (13–16).

In this work a decellularization pipeline is proposed, to obtain a safe and highly preserved decellularized small intestine (dSI). For this, penetration enhancers will be used as a strategy to enhance the decellularization efficiency while reducing the time of exposure to detergents. Parameters such as decellularization efficiency, biochemical composition, mechanical integrity and cytocompatibility have been assessed at different steps of the methodology to achieve an optimized tissue-specific protocol for small intestine decellularization. Also, the most adequate side (serosa or submucosa surface) to contact the wound will be investigated.

## 2. Materials and Methods

### 2.1. Intestine harvesting

Porcine small intestine from three freshly slaughter 6-months pigs, were harvested and transported to the facilities of *Universidade Católica Portuguesa* from Seara S.A. company, Vila Nova de Famalicão, Portugal, properly conditioned and refrigerated in a cold box on ice to prevent tissue degradation. The small intestine was cleaned with cold 1XPBS, the superficial fat was removed using a scalp and tweezers, and then the intestine was divided in 5 cm portions and stored at −20°C to be randomly assigned to untreated and treated tissue to minimize variability.

### 2.2. Decellularization

The cell content of the material was removed from the tissue by immersion and agitation as represented in Figure 1.

**Figure 1.**
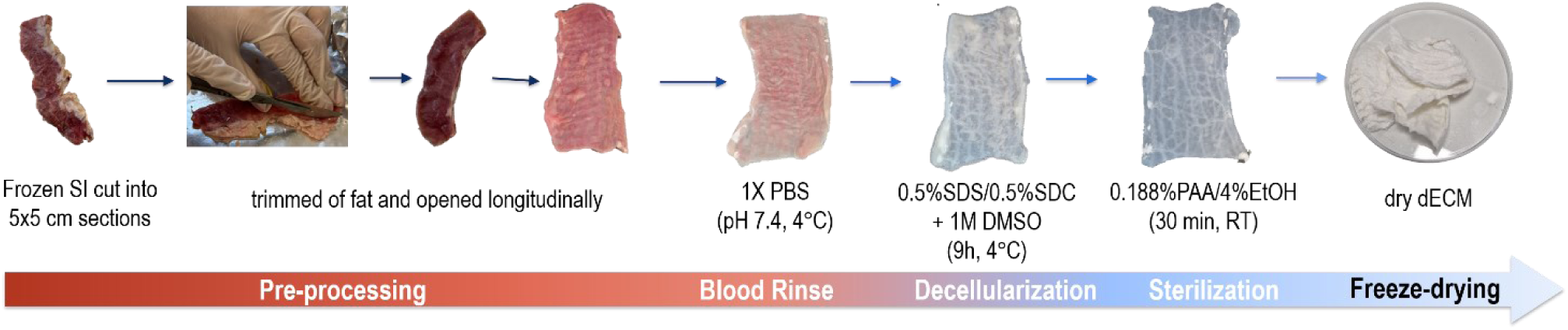
Overview of the key steps involved in small intestine decellularization.

Briefly, intestine samples were open longitudinal, washed with 1X PBS (pH 7.4, 4°C) with chloramphenicol (CLO, 0.06 mg/mL) until the solution ran clear (30 min, approximal 5 cycles). After the tissues are incubated in 0.5% sodium deoxycholate (SDC, Sigma-Aldrich, Missouri, United States) and 0.5% sodium dodecyl sulfate (SDS, Sigma-Aldrich, Missouri, United States) in 1M DMSO (Honeywell, Charlotte, NC, USA) for two 4h30min cycles, separated by overnight washes in distilled water with chloramphenicol (CLO, 0.06 mg/mL). DMSO was used based on its well-characterized ability to alter the physical structure of lipid membranes, that shown DMSO increases membrane fluidity and induces transient pore formation at higher concentrations, facilitating membrane disruption and cell removal (17).

The tissue underwent sterilization by immersion in a solution of 0.188% peracetic acid in 4% ethanol (Thermo scientific, Waltham, MA, USA) for 30 minutes at room temperature followed by rinsing in 1X PBS. Samples were washed for a total of 72h with exchanges every 2h a day until left overnight. Lastly, sterilization was carried out using a peracetic acid/ethanol solution (0.188% PAA / 4.8% EtOH) for 30 min under agitation (100 rpm, RT). Samples were washed in PBS for 5 cycles to ensure removal of sterilizing agents followed by ultrapure water rinses for a 24h cycle. The resulting tissue was frozen at −80°C, lyophilized for 72 h and keep at −80°C until further use. All steps were performed while the specimens were kept in a cold box on ice to prevent protease activity under agitation excepted clearly stated otherwise.

### 2.3. Scanning electron microscopy (SEM)

Samples (n=3) were crosslinked with 2.5% glutaraldehyde (Sigma-Aldrich, US) in 1X PB buffer (pH 7.4), overnight at 4°C. The samples were cleaned 5 times in 1X PB buffer (pH 7.4) for 1-2 min each. After fixing, the specimens were dehydrated in gradually increasing EtOH rinses (10%, 25%, 50%, 75%, and 95%) for 1 h each and finally rinsed twice in 99.8% ETOH (Honeywell, US) for 10 min per wash to fully dehydrate the specimens. Ethanol was removed by critical point drying. Cross-section samples were cut using liquid nitrogen before fixation All samples were coated with Au/Pd with the SC7640 sputter coater (Polaron, UK) at 20 mV prior examination on a Phenom™ Pro Desktop scanning electron microscope (SEM) (Thermo Scientific™, US). Visualization was performed at 15 kV and digital images were acquired at 500x and 5000x magnification.

### 2.4. Histology

Small intestine samples were fixed in neutral-buffered formalin (10%) for 24h – 48h, dehydrated, and embedded in paraffin. Tissue sections (3 μm) were cut and stained with hematoxylin and eosin (H&E), Verhoeff, Masson’s Trichrome and Alcian blue. The prepared samples were evaluated under a Nikon Eclipse E50i light microscope (Nikon Corporation Instruments Company, Tokyo, Japan) and were analyzed using the ImageJ® software.

### 2.5. Biochemical analysis

#### 2.5.1 DNA quantification

DNA content was analysed from native and decellularized samples (n = 5) to assess decellularization efficiency. Tissue was freeze dried and minced for DNA quantification using Genomic DNA Kit Tissue (GRISP, Portugal), according to the manufacturer’s instructions with a few modifications. Sample digestion period was extended until no visible ECM particles weren’t observed. Purified DNA was then quantified using Quant-iT™ PicoGreen™ following the manufacturer’s instructions. DNA yield was then measured using the BioTek Synergy H1 microplate reader (Agilent Technologies, Inc., Santa Clara, California, USA) by fluorescence excitation at 502 nm and emission of 523–524 nm.

#### 2.5.2. Elastin, sulphated glycosaminoglycans and soluble collagen

Alfa elastin, sulfated glycosaminoglycans (sGAGs), and soluble collagen content in the native and decellularized scaffolds (n=5) were quantified using the Fastin™ Elastin Assay Kit (biocolor, United Kingdom), Blyscan™ sGAG Quantitative Dye-Binding Assay Kit (n=5) (biocolor, United Kingdom), and Sircol™ Soluble Collagen Assay kit (n=5) (biocolor, United Kingdom), respectively, following the manufacturers’ instructions.

#### 2.5.3. Lipid quantification

Total lipid (n=5) content was determined through the chloroform method with some modifications (18). Briefly, to 50 mg of milled tissue it was added chloroform (Honeywell, Wabash, IN, USA), methanol (Honeywell, Wabash, IN, USA), and 50 mM potassium phosphate buffer (Sigma-Aldrich, St. Louis, MO, USA), followed by incubation under agitation (room temperature, 2h). Afterwards, chloroform and 50 mM potassium phosphate buffer were added, mixed, and then left to rest until layers formed within the solution. The chloroform layer was then transferred to a dry glass tube and left to volatilized in a water bath at 60 °C to get the dry extracted lipid content. The total lipid content (LC) was calculated as follows:

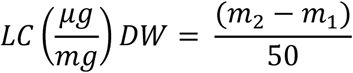

### 2.6. Residual SDS

Residual SDS (n = 3) was quantified using an SDS Detection and Estimation Kit (G Biosciences, St. Louis, MO) according to the manufacturer’s instructions on milled tissue. Briefly, 1 mL of methylene blue solution was combined with 0.5 mL of extraction buffer and 20 mg of the milled tissue containing SDS, followed by vortex mixing for 30 s. Subsequently, 1 mL of chloroform was added and the mixture was vortexed for an additional 30 s. After phase separation for 5 min, the lower chloroform layer was collected for absorbance measurement at 600 nm. SDS concentrations were determined using a calibration curve generated from known standards.

### 2.7. Swelling ability (SA)

Swelling test (n = 7) was conducted to determine water holding capacity of the native and decellularized tissue. Dry weight (W_0_) of the specimen was calculated. Sample was immersed into 40 mL of 1X PBS or SWF A (19) (pH 7.4) at 37 °C (W_s_) was register after 1h, 2h,4h, 8h 12h, 24h and after every 24h until plateau. The swelling ratio was calculated using the equation below:

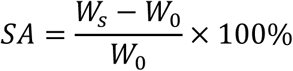

### 2.8. Fourier Transform Infrared (FTIR) Spectroscopy

Mid-infrared spectra of the intestine samples (n=3) were obtained on a Fourier transform PerkinElmer Spectrum BX FTIR System spectrophotometer (USA) with a DTGS detector. Spectra were acquired in diffuse reflectance mode through a PIKE Technologies Gladi attenuated total reflectance (ATR) accessory within the wavenumber interval of 4000 to 600 cm^-1^, with a resolution of 4 cm^-1^. Each spectrum resulted from 32 scan co-additions. Samples were placed in the ATR crystal and a constant pressure was applied. The ATR crystal was cleaned, and a background was acquired between each sample. Three spectra per sample were acquired. Infrared spectra were modelled through a principal component analysis (PCA) (20). Prior modelling spectra were pre-processed with standard normal variate (SNV) (21) and the Savitzky-Golay filter (15 smoothing points, 2^nd^ order polynomial and 2^nd^ derivative) (22) and further mean centered. Spectral pre-processing and modelling were performed in MATLAB R2023a (MathWorks, Natick, MA) and PLS Toolbox 9.2.1 (Eigenvector Research, Manson, WA).

### 2.9. Uniaxial tensile test

Mechanical properties (n=8) of treated and native tissue were performed according to Ferrara *et al.* with a few modifications (23). The specimens were longitudinally cut using scalpel and a 3D printed dumbbell mold with dimensions according to ASTM D412 was used to favor rupture inside the fracture area (Figure 2A-C) (24). The mold was placed circumferentially. Samples were immersed in 1X PBS (pH 7.4) supplemented with 0.02% (w/v) sodium azide at 37°C for 24h before testing, to ensure sample hydration. Mechanical tests were performed using a texturometer (TA.XT PLUS, Texture Analyzer, UK) by analyzing the degree of extension of the specimen. The stretching was performed using mini tensile grips (Stable Micro Systems). The system is equipped with a 5kg load-cell anchored to the crosshead and by two grips. The mechanical test consisted of two analysis phases, preconditioning to minimize variations exhibited by soft tissue followed by uniaxial extension: (a) preconditioning phase consisting of 10 cycles of loading-unloading at constant crossed speed of 1 mm/min (Figure 2D) in the elastic zone followed by (b) uniaxial tensile extension using constant crossed speed of 1 mm/min until specimen failure. During testing sample frequency was 50 Hz. Test were considered valid when sample fractured inside the rupture area The results were obtained by analysing the obtained stress-strain curve to obtain maximum elongation, elastic modulus and ultimate strength.

**Figure 2.**
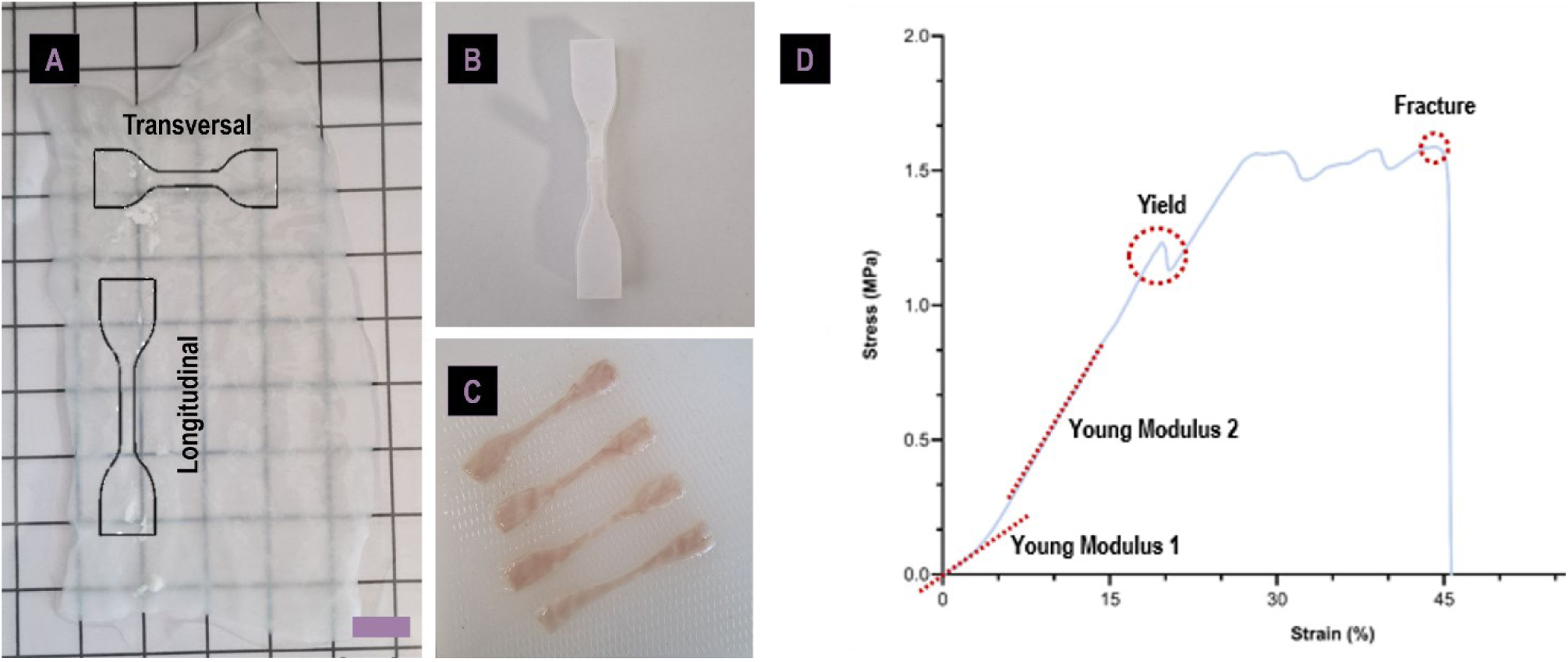
(A) Representation of the direction of the small intestine specimens for uniaxial testing. (B) 3d printed dumbed mold according to ASTM D412 and (C) prepared native small intestine specimens. (D) Representative cyclic pre-conditioning stress–strain curve of the tested tissue.

### 2.10. Thickness

Micrometre Adamel Lhomargy MI 21 was used to measure sample thickness (n=11). Briefly, samples were immersed in 1X PBS (pH 7.4) supplemented with 0.02% (w/v) sodium azide at 37°C for 24h before testing, to ensure sample hydration. Each sample of about 10×5 cm was measured into 10 equally space points and the value average to considered the sample thickness according to the ASTM D6988.

### 2.11. Water vapor transmission rate (WVTR)

Water vapor transmission rate was measured by gravimetric method placing freeze dried native and treated small intestine membranes (n=3) in sealed metal support with paraffin and pressure. Briefly, water was placed inside the metal support and the system was weighed at different periods from 0h to 190h. Samples were kept in a room with control vapor saturation pressure (52%) and temperature (23°C) according to the ASTM E96-2005

### 2.12. Differential scanning calorimetry (DSC)

Thermal stability of the native and decellularized ECM (n=3) was assessed using a DSC 204 F1 Phoenix^®^ (NETZSCH, Selb, Bayern, Germany). The samples, which weighed between 2 and 3 mg, were heated at temperatures ranging from 0 to 280°C. The process was conducted at a rate of 10°C/min using a nitrogen flow of 100 mL/min to serve as a protective atmosphere during the entire assay.

## 3. Microbial invasion

To assess the barrier ability of tissue scaffolds and *in vitro* bacterial invasion assay was performed. Briefly, 3 sterile glass vials were filled with 1 mL of lysogeny broth (LB). Vials were sealed with parafilm, the tissue scaffolds and two commercial band-aids, and a last vial using perforated parafilm to serve as disrupted barrier condition. All samples were maintained at 37°C and after 1, 3, 7 and 14 days bacterial growth within the broth was evaluated by measuring the optical density of the broth at 600 nm. Sterile media was used as blank.

## 4. Degradation behavior

A degradation assay (n = 9) was performed using a gravimetric approach with the addition of collagenase from *Clostridium histolyticum* (CCH, type I, ≥ 125 CDU/ mg solid, Sigma-Aldrich) and hydrogen peroxide (30%, Sigma-Aldrich) at different pH values to simulate distinct burn-wound environments (25). Briefly, tissues were first allowed to swell for 72 h in simulated wound fluid A (SWF A) (19). After swelling, samples were gently blotted dry with sterile filter paper and weighed to obtain the initial wet weight. Samples were then transferred to one of five degradation solutions. The control solution consisted of SWF A at pH 7.4 and served as the base formulation for all other conditions. The remaining solutions contained either collagenase alone (0.08 µg/mL or 0.33 µg/mL) or collagenase combined with H₂O₂ (50 mM or 166 mM), adjusted to pH 6.0 or pH 8.0. The conditions are summarized in table 1(26).

**Table 1.** Composition of degradation medias.

| Label | Composition | pH |
| --- | --- | --- |
| SWF A | SWF A | 7.4 |
| SWF A w/CCH (pH 6.0) | 0.08 ug/mL CCH | 6.0 |
| SWF A w/CCH and H<sub>2</sub>O<sub>2</sub> (pH 6.0) | 0.08 ug/mL CCH and 50 mM H<sub>2</sub>O<sub>2</sub> | 6.0 |
| SWF A w/CCH (pH 8.0) | 0.33 ug/mL CCH | 8.0 |
| SWF A w/CCH and H<sub>2</sub>O<sub>2</sub> (pH 8.0) | 0.33 ug/mL CCH and 166 mM H<sub>2</sub>O<sub>2</sub> | 8.0 |

Collagenase levels were chosen based on two criteria. First staying bellow commonly used 2.5 µg/mg of CCH used since systematic review, reported this *in vitro* collagenase concentrations are generally higher than physiological levels and overestimate degradation rates leading to large gaps between *in vitro* and *in vivo* (*27*). The range was further defined considering the MMP-8 and MMP-9 cumulative values find in burn wound exudates (28). H₂O₂ levels were selected to span from mildly to clearly damaging conditions. In a murine wound model, 10 mM H₂O₂ accelerated closure, whereas 166 mM delayed healing, reduced connective tissue formation (29). Accordingly, a high H₂O₂ condition (∼166 mM) reflects a documented delayed-healing environment, while a lower dose (e.g., 50 mM) represents sub-maximal oxidative stress within the damaging range (29–31).

Samples were weighted at 3 day intervals throughout a 26-day period. At each timepoint, the samples were gently blotted to remove excess solution and weighed. The incubation solution was refreshed after each measurement. Mass loss was calculated using the following equation:

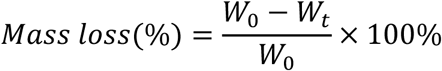

where W_0_ is the initial weight after 72h sweeling and W_t_ the dry weight at a given timepoint.

## 5. *In vitro* dECM surface screening by direct cell seeding

Human dermal fibroblasts (HDF, passages 15–20) were cultured in high-glucose DMEM supplemented with 10% fetal bovine serum (FBS). HaCaT keratinocytes (passages 28–32) were cultured in high-glucose DMEM without calcium, supplemented with 10% FBS and 30 µM calcium chloride. All cells were maintained at 37 °C in a humidified atmosphere containing 5% CO₂. Human dermal fibroblasts (HDFs) and HaCaT keratinocytes were seeded onto the decellularized extracellular matrix scaffolds both in the serosa and submucosal sides. Circular scaffolds (6 mm in diameter) were prepared using a sterile Kai disposable dermal biopsy punch and placed individually in well plates prior to cell seeding. For each scaffold, a concentrated cell suspension was prepared to deliver 10,000 cells in a final volume of 10 µL. This suspension was carefully pipetted directly onto the surface of the dECM to promote localized cell attachment and limit cell loss. Following seeding, constructs were incubated under standard culture conditions for 4 h to allow initial cell adhesion during this time and every 2 h, 20 µL of culture medium was gently dispensed at the base of the well, near but not directly onto the scaffold, to maintain humidity and prevent scaffold dehydration without disturbing the cell droplet. After the 4 h attachment phase, an appropriate volume of complete culture medium was added to each well to fully submerge the scaffolds, and cultures were maintained under standard conditions for subsequent experiments.

### 5.1. Metabolic activity

The metabolic activity (n=3) of cells was quantified by performing a PrestoBlue assay. After each time-point, a 10% (v/v) presto blue working solution was prepared in standard culture medium, added to samples and left for 2 hours at 37 °C and 5% CO2 incubator. Supernatant fluorescence was recorded at an excitation wavelength of 560 nm and an emission wavelength of 590 nm using a CLARIOstar Plus (BMG LABTECH, Ortenberg, Germany) microplate reader. Presto blue working solution was used as a blank

### 5.1. Proliferation

Cellular proliferation was assessed by BrdU (colorimetric) incorporation assay (n=3) using a ELISA kit (Roche, Basel, Switzerland) was performed according to the manufacturer’s instructions. Briefly, after the 4 hours incubation in the cell seeding, 200 µL of standard culture medium supplemented with a BrdU labeling solution was added. Media was refreshed every 3 days and new labeling solution was added to access cumulative proliferation. After each time-point, samples were fixed and quantification followed the manufacturer’ s instructions. Briefly, at each time point, samples were fixed in oven at 60°C for 1h followed by DNA denaturation with the denaturation solution provided, followed by incubation with a peroxidase-conjugated anti-BrdU antibody. After washing, substrate solution was added for 30 min. Cell free scaffolds were used as blanks and absorbance was measured using CLARIOstar Plus (BMG LABTECH, Ortenberg, Germany) microplate reader at 650 nm. Samples without seeded cells were used as blank control and a total of 3 independent samples were tested.

## 6. Statistical Analysis

Statistical analyses were carried using GraphPad Prism. Data was assessed for normality using the Shapiro-Wilk test. One-way analysis of variance (ANOVA) was conducted for normally distributed data, followed by Tukey’s post hoc test to identify pairwise differences. For non-normally distributed data, the Kruskal-Walli’s test was applied. Results are presented as mean ± standard deviation (SD). Differences between data sets were considered statistically significant if *p* < 0.05.

## 3. Results

### 3.1. Assessment of Decellularization Outcomes

The macroscopic and H&E histological images of the porcine small intestine tissue at each stage of the decellularization process is shown in Figure 3.

**Figure 3.**
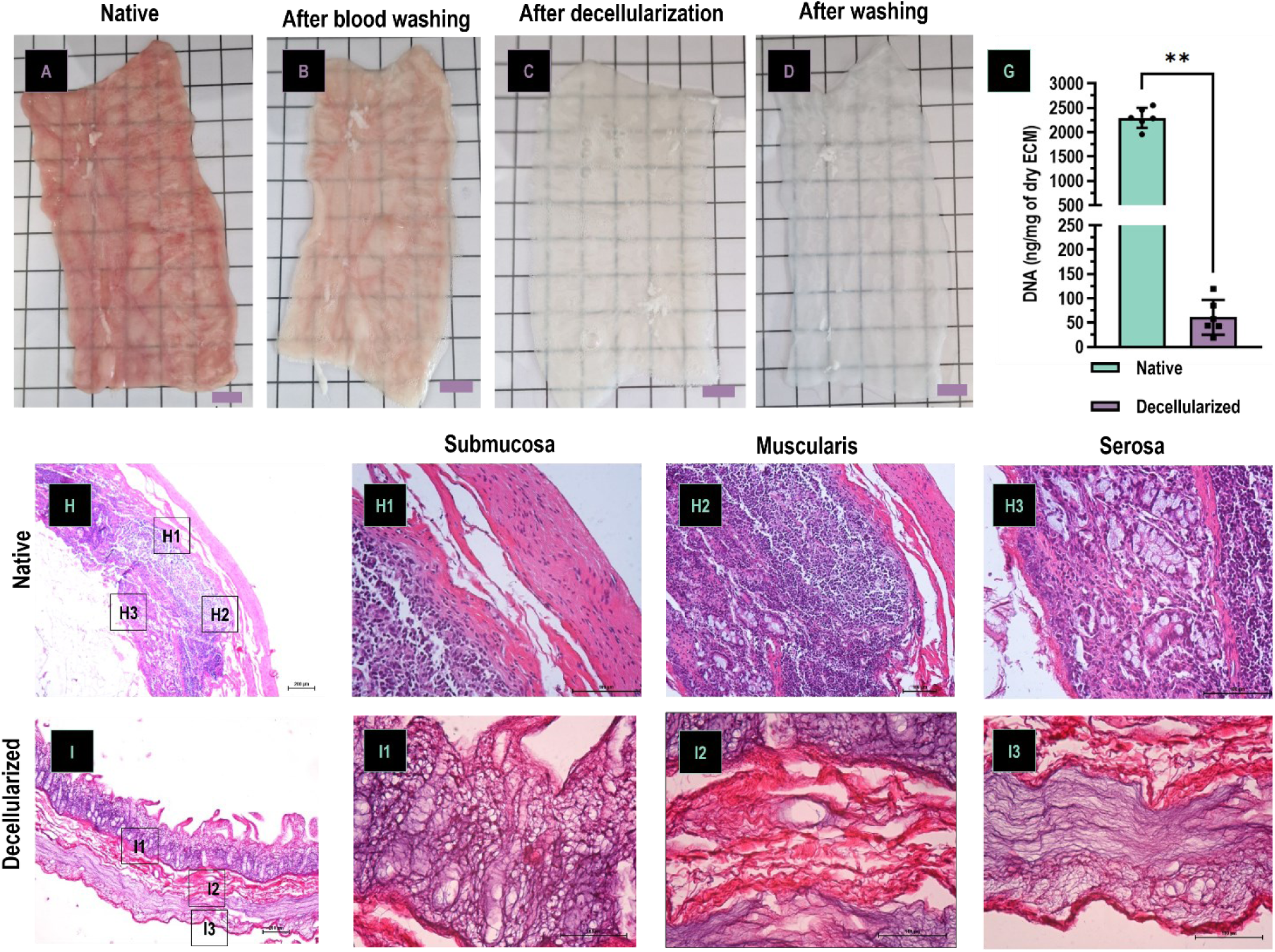
Macroscopic images of the small intestine samples and decellularization assessment: (A) native small intestine, (B) after blood washing with PBS, (C) after detergent decellularization stage, and (D) after washing and sterilization. (G) DNA quantification of native and treated tissue. H&E staining of small intestine samples: (H1-3) native small intestine and (I1-3) after decellularization protocol. Data are presented as mean ± standard deviation, with significant differences indicated (*p < 0.05).

While native tissue exhibited a natural pinkish coloration the treated segments displayed a whitish appearance after decellularization, retained a similar macrostructural appearance after the washing step with a clearer and more translucid surface. To evaluate the efficiency of the decellularization protocols, histological analysis using H&E staining and DNA quantification were performed on the tissues. The native small intestine (Figure 3 H1-H3) exhibited abundant nuclei, indicative of the presence of cellular content. Following decellularization (Figure 3 I1-I3) dECM demonstrated extensive removal of cellular material, with occasional residual nuclei visible in certain regions. Moreover, the submucosa layer preserved villosities structure, while the tunica serosa ECM structure appeared to have well organized fibers along the longitudinal axis. This suggests that the decellularization method implemented was effective in removing cellular content, without compromising ECM architecture.

Quantitative analysis of residual DNA (Figure 3 G) revealed significant DNA reduction in samples treated with the proposed protocol compared to native tissue. The native intestine exhibited a DNA content of approximately 2296 ± 206 ng/mg of dry ECM, whereas the decellularization protocol presented DNA levels of approximately 49.71 ± 24.08 ng/mg, which represents a statistically significant reduction (**, p < 0.01). These findings confirm that the decellularization protocol implemented is effective to decellularize small intestine tissue while allowing for preservation of ECM architecture, highlighting its potential as an advanced decellularization approach. Moreover, SDS a cytotoxic component used in the decellularization protocol implemented was quantified after 24h, 48h and 72h of washing and it was found to be at 0.00088% ± 0.0002% after the first 24h washing cycle, i.e. below the levels considered cytotoxic of 0.001% (32). This component was established has the metric for harsh component removal has it is reported to be challenging to remove after extension tissue immersion due to established protein-SDS complex formation (33).

### 3.2. Tissue Microstructure and properties

SEM was performed on the tunica serosa, adventitia and in a cross section along the circumferential direction of the native and decellularized small intestine segments to examine the morphology and structure. As shown in Figure 4, in the intact native small intestine, a dense network of fibers is present in the tunica serosa (Figure 4 A1). After the decellularization treatment, the ECM network of fibers presents a well-organized structure (Figure 4 B1). Moreover, images in Figure 4 B1 also reveal the tape shaped fibrillar network that was retained after treatment in the tunica serosa. In the tunica submucosa of the native small intestine, a dense structure of debris is present in the ECM network (Figure 4 A2), likely the result of cells, mucus and bacteria naturally present in small intestine. After the decellularization protocol, the ECM structure is clearly revealed with the removal of cells and debris. This structure presents a well-preserved villus-crypt architecture and porosity along the villous. Cross-section images in Figure 4 A3 show the three layers of the small intestine with distinct ECM structural organization. After treatment the individual layers are still identifiable with distinct organization and an intact boundary between each tunica’s, however the tissue’s thickness appeared lower has compared to native tissue (Figure 4 G). Native tissue presented a 0.50 ± 0.063 µm while after decellularization the value decreases to 0.40 ± 0.050 µm with significant difference in the mean thickness (p=0.0006). Water vapor transmission rate and swelling assays were performed to understand the water vapor and water uptake capabilities of the material as dressing for burn wounds.

**Figure 4.**
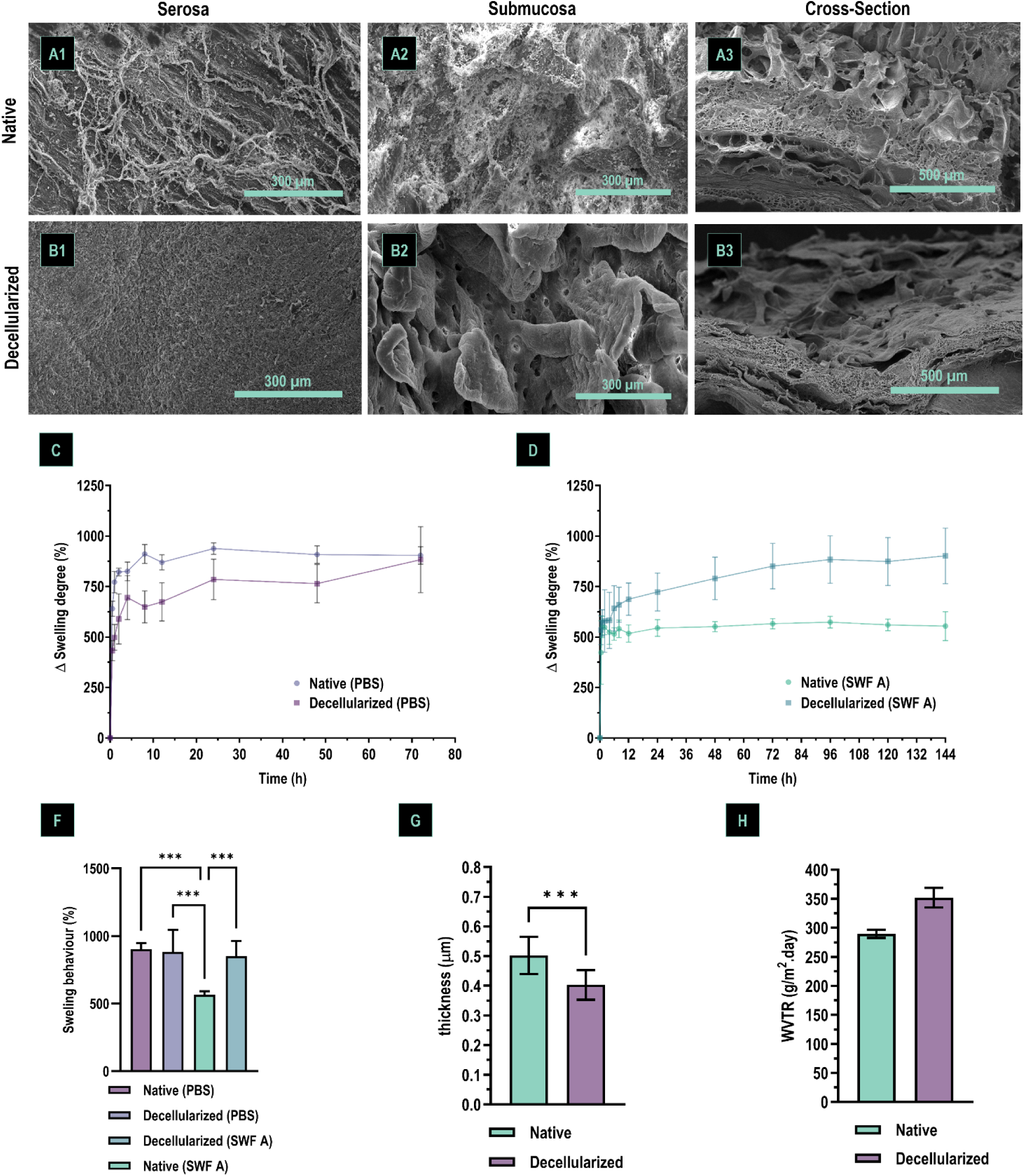
Scanning electron micrographs of small intestine: (A1–A3) native tissue and (B1–B3) tissue after decellularization. Cross-section images made in a circumferential direction. (C) Swelling behaviour of small intestine in PBS and (D) simulated wound fluid. (F) Maximum swelling of native and treated small intestine, (G) Material thickness before and after treatment and (H) water vapor transmission rate of small intestine. Data are presented as mean ± SD.

The mean WVTR of the native small intestine was 289.57 g/m^2^.day and after treatment there was a slight increase in this rate to 351.88, without statistically significant differences between the groups. All groups exhibited an initial rapid increase in swelling within the first 2h30 min, followed by a plateau phase after approximately 10 hours. The native small intestine demonstrated a similar equilibrium swelling percentage (904% ± 43.33%) in PBS as compared to the treated tissue (883% ± 163.35%), indicating preserved sweeling capability after 72h. The decellularization methods did not impacted the tissue’s hydrophilic properties with no statistically significant differences among the groups by the end of the swelling assay.

In contrast, tissue swelling in SWF required a longer time to reach a plateau than in PBS and showed significant differences between native and decellularized tissue, respectively (565.63% ± 25.36%, 851% ± 113%, p=0.0006).

### 3.3. Biochemical Composition and Histological Architecture

Histological analysis and biochemical assay results are summarized in Figure 5. As determined by Verhoeff staining, the ECM red stain elastic fiber content was observed in the dECM group with a more disorganized and coiled fiber arrangement in the muscularis layer and a well-defined orientation along the longitudinal axis in the serosa layer (upper panels in Figure 5 A1 to 3 and B1 to 3) than the native tissue. However, alpha elastin quantification revealed a statistically significant decrease in content of approximal 57.7% (p=0.0233). MT staining was also conducted to examine the ultrastructural distribution and quality of the collagen content. In the treated small intestine group, collagen was observed along the three layers of the tissue, marked by the blue stain, (Figure 5 C1 to 5 C3), mainly in the muscularis layer comparable to the elastin content, with similar fiber arrangements as described.

**Figure 5.**
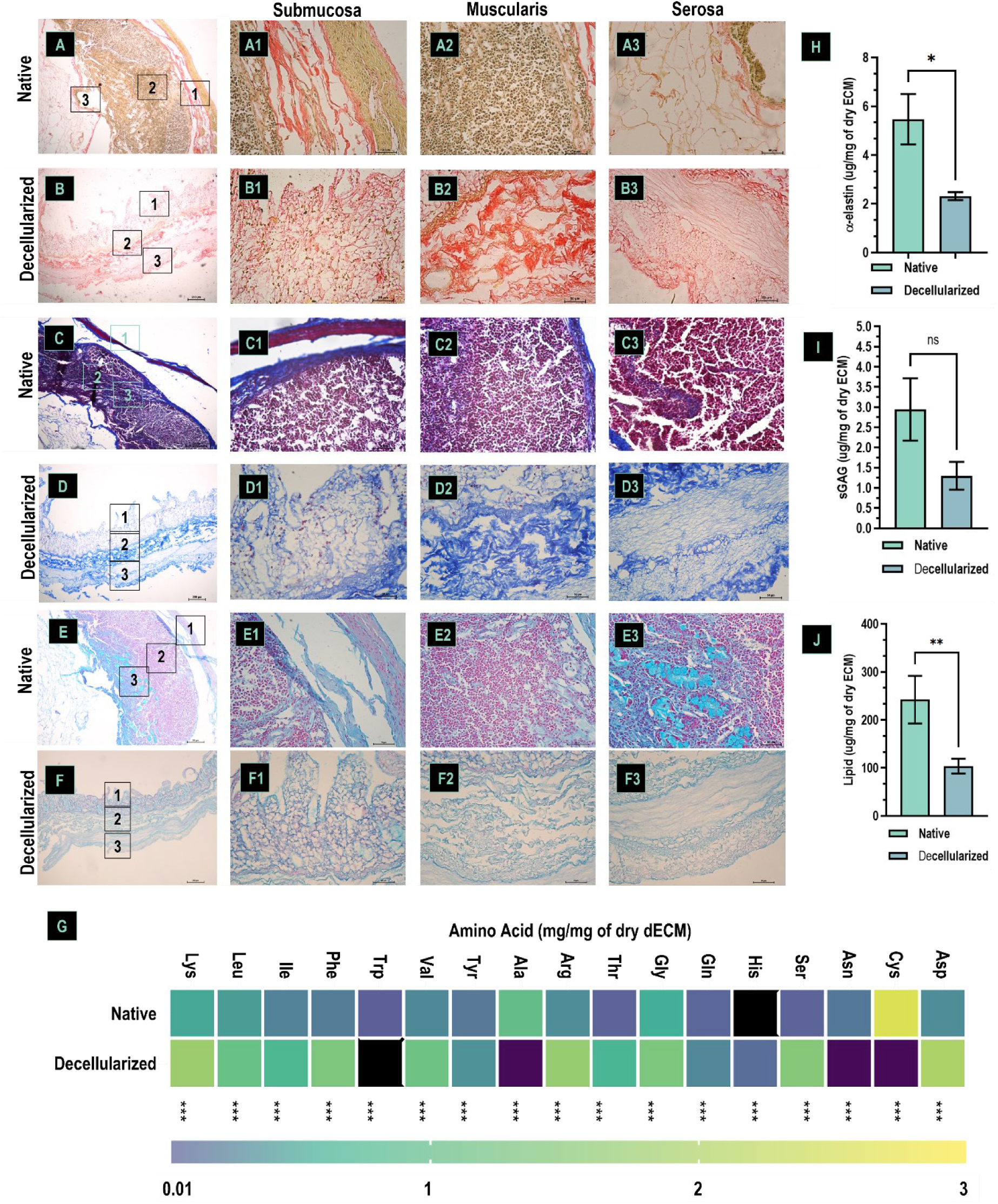
Histological and biochemical assays of porcine small intestine samples: Verhoeff stain of (A1-3) native small intestine and (B1-3) after decellularization protocol, Masson trichomic stain of (C1-3) native small intestine and (D1-3) after decellularization protocol and alcian blue stain of (E1-3) native small intestine and (F1-3) after decellularization protocol. (H) Alpha elastin, (I) sulphated GAGs, (J) lipid of native and decellularized tissue. (G) Aminoacidic profile of native and decellularized tissue, where the black square marks outside of the lower limit of detection and dark purple bigger than 3. Black bar-native small intestine and blue bar decellularized small intestine. Data are presented as mean ± standard deviation, with significant differences indicated (*p < 0.05 and **p<0.005).

Alcian Blue staining was performed to reveal the GAG content of the small intestine matrix, an essential component of the ECM. The stain showed that the GAG content was present before and after decellularization, mainly on the serosa layer (Figure 5 E1-3 and F1-3). Similar trends were observed upon the quantitative analysis of the GAG content shown in Figure 5 I, with no statistically significant differences between native and treated tissue. Different trend was observed for lipid content (Figure 6 L), which significantly decreased by 49.1% (p=0.0087) after decellularization compared with native small intestine.

**Figure 6.**
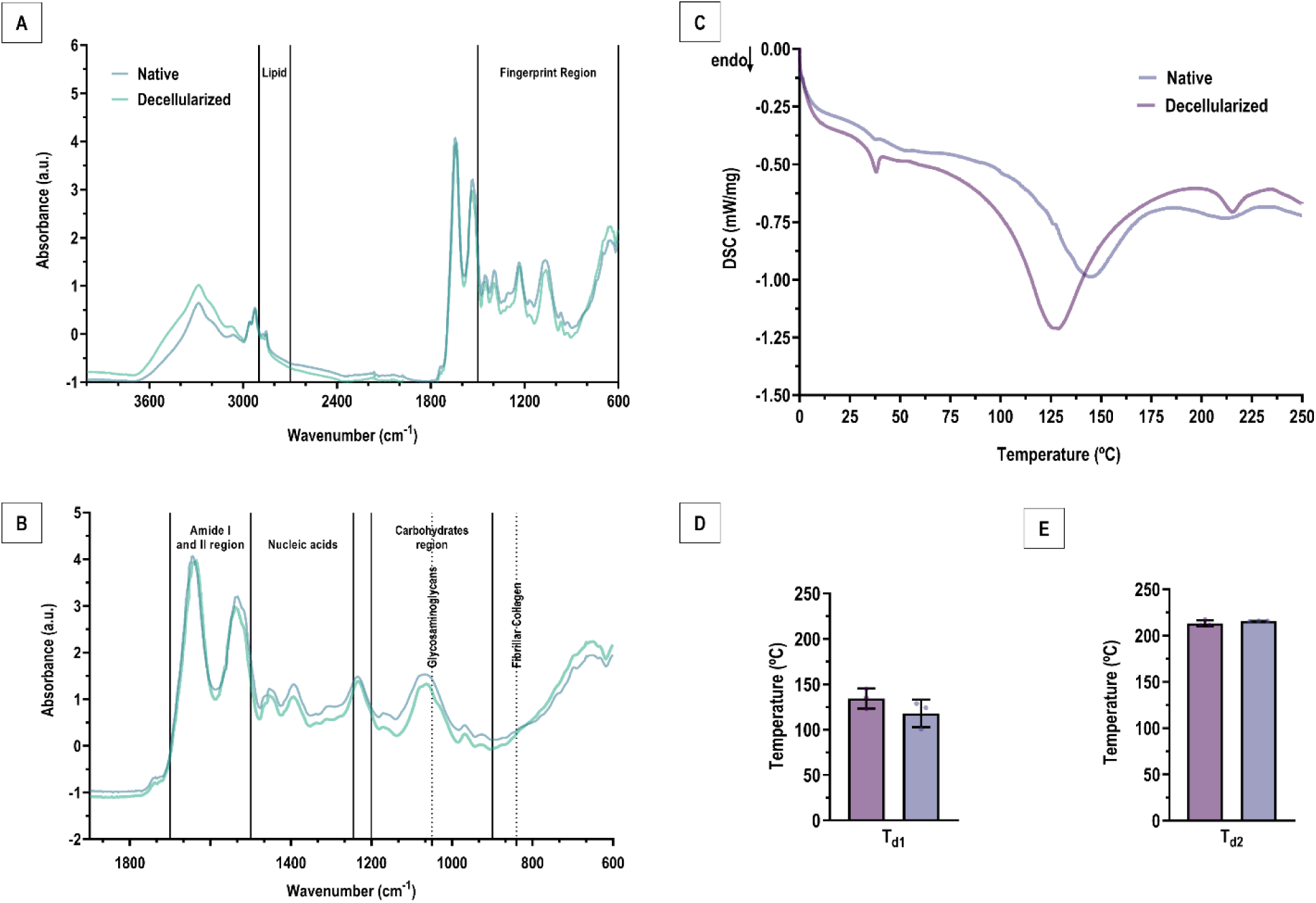
Mid-infrared (A, B) mean spectra of the intestine samples. DSC thermogram (C), and temperature peaks (Td) (D, E) during collagen fiber denaturation of native and treated small intestine. Blue line-native small intestine and green line decellularized small intestine.

The amino acid profile in native and decellularized tissue is presented in Figure 5 G. Many amino acids (e.g., Cys, Ala, Gly, Arg, Lys) were substantially higher in the decellularized tissue than in the native tissue. Some of aminoacid residues were reduced or nearly absent in the decellularized tissue (e.g., Asn, His), or absent (e.g. Trp).

The FTIR-ATR analysis was performed to investigate the chemical structure of the native and decellularized intestine ECM (Figure 6 A). Intestine samples infrared spectra exhibit the typical bands assigned to lipids (2900-2700 cm^-1^), proteins (amide I and amide II region: 1700-1500 cm^-1^), nucleic acids (1500-1200 cm^-1^) and carbohydrates (1200-900 cm^-1^) (Figure 6 B). Decellularized dECM spectra (green line) present a clear decrease of the infrared bands attributed to lipid vibrations, suggesting that the decellularization process resulted in lipid removal. Also, the broad bands attributed to nucleic acid vibrations decrease in the region from 1500-1200cm^-1^, pointing to a lower content of these molecules in the decellularized material. On the other hand, it is possible to clearly notice the presence of collagen after the decellularization process (black lines) due to its typical vibration bands (amide I, II and three weaker bands centred at 1245 cm^-1^) and the appearance of a new band nearly at 840 cm^-1^ typical of fibrillar collagen (Figure 6 B). The presence of GAGs after the decellularization can also be proved through their typical infrared vibration bands, namely: the increase in the broad band centered at 1050 cm-1 (typical of the symmetric stretching of C-C-C; C-C-O, C-O-C and SO_3_^-^ of GAGs).

Native small intestine started the endothermic process at approximately 134.4 ± 11.1°C (Figure 6 C) and had denaturation temperature peaks (Td) at approximately 213.2 ± 3.4°C (Figure 6 D to 6 E). For the dECM the endothermic process began at approximately 117.8 ± 15.1°C (Figure 6 C), and its denaturation temperature was the first temperature peak was slightly higher (approximately 215.7 ± 0.3°C) compared with that of the native tissue (Figure 6 D to 6 E). There were no statistically significant differences between the groups.

### 3.4. Mechanical Properties Under Tensile Loading

To evaluate the effects of the applied treatment on the mechanical material properties uniaxial tensile assay were performed on the tissues. The stress-strain curves presented in Figures 7 A and 7 L highlight the mechanical responses of the samples under tensile testing. In transversal orientations, tissues consistently exhibited higher extensibility compared to longitudinal orientation, while in the longitudinal direction tissues presented higher stress resistance, reflecting the anisotropic nature of the small intestine.

**Figure 7.**
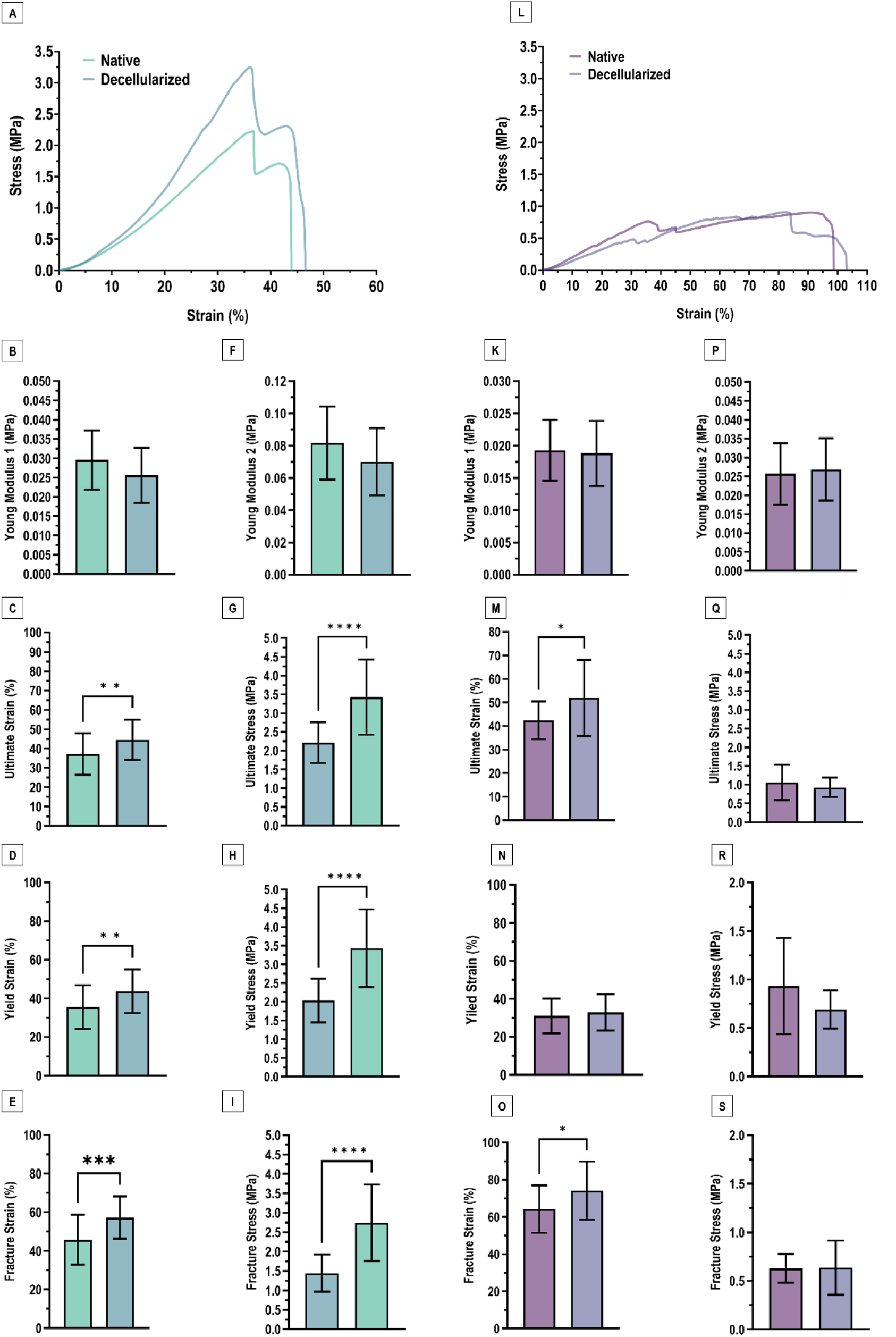
Uniaxial tensile mechanical properties of the native and treated small intestine. Representative (A,L) uniaxial tensile break stress–strain curve of the tested tissues in a longitudinal and transversal direction respectively. (B, K) Young’s modulus from the toe region (Young’s modulus 1; B, K) and high-stress region (Young’s modulus 2; F, P) of the stress–strain curve. (G, Q) Ultimate, (H, R) yield and (I, S) fracture stress, and (C, M) Ultimate, (D, N) yield and (E, O) fracture strain of the tested tissues. Each condition contained *n* = 17. Data are presented as mean ± standard deviation, with significant differences indicated (*p < 0.05).

The Young’s modulus was calculated from two distinct regions of the stress-strain curve, provided insights into tissue changes of its nonlinear response with increasing stiffness. Figures 7 B, 7 F, 7 K and 7 P illustrates the Young’s Modulus taken at low and high stress areas. The longitudinal orientation produces an overall higher elastic modulus than the transversal direction. The mean Young’s modulus 1 of the native tissue in the transversal direction was 0.019 ± 0.005 MPa while in the longitudinal direction was 0.030 ± 0.008 MPa. For the treated tissue, the mean elastic modulus, was 0.019 ± 0.005 MPa and 0.026 ± 0.007 MPa for the transversal and longitudinal direction, respectively (Figure 7 B, F, K and P). Interestingly, the treated tissue resulted in Young’s Modulus 1 values comparable to the native protocol in both orientations, with no significant differences with the native tissue.

Figure 7 F and 7 P present Young’s Modulus 2, representing the high stress region of the stress-strain curve. In this region, the tissues again demonstrated higher stiffness in the transversal orientation. The mean young modulus 2 of the native tissue was 0.026 ± 0.008 MPa in the transversal direction and 0.082 ± 0.023 MPa in the longitudinal one; while, for the treated tissue, the mean elastic modulus, was 0.027 ± 0.008 MPa and 0.07 ± 0.02 MPa, respectively (Figure 7 F and 7 P). In this region, the tissues again demonstrated higher stiffness in the longitudinal orientation. However, the differences between native and decellularized groups were not statistically significant for either orientation.

Figure 7 G, 7 H, 7 I and 7 Q, 7 R, 7 S show the ultimate, yield, and fracture stress, while the matching strain parameters appear in Figure 7 C, 7 D, 7 E and 7 M, 7 N, 7 O. Looking at the longitudinal orientation, decellularized tissues lose stress-bearing strength compared to native tissue. Peak, yield, and fracture stress all decrease after decellularization (Figure 7 G to 7 I), which shows that decellularization weakens longitudinal tensile strength. On the other hand, when looking at the transversal orientation, this difference is less pronounced. Stress values stay relatively similar between native and decellularized groups as well as strain results. In the longitudinal direction, decellularized tissues behave differently under ultimate, yield, and fracture strain which all shift compared to native controls (Figure 7 M to 7 O). But in the transversal direction, strain values scarcely change after decellularization. In short, decellularization clearly affects tensile strength and how the tissue fails in the longitudinal direction but leaves the transversal mechanical properties mostly preserved. This directional effect points to the tissue’s anisotropic structure and suggests that the matrix components responsible for handling load along the longitudinal axis are predominately affected by the decellularization.

To further investigate the tissue treatment over the dynamic mechanical properties a cyclic uniaxial tensile test on the tissue was performed, stretching it in both the longitudinal and transverse directions. Figure 8 A to D displays representative stress–strain curves for both native and decellularized samples. In every case, the tissue exhibited a distinct hysteresis loop between loading and unloading, a typical indication of dissipating energy and viscoelastic behaviour. Consistent with the results from tensile tests until break, tissues in the longitudinal direction withstood greater stress. Figure 8 E and G illustrate the variation in peak stress with each cycle at a constant strain. Regardless of the group, peak stress decreased over time as the number of cycles increased, indicating tissue relaxation and adaptation to repeated stretching. Notably, native tissues exhibited a more rapid decline in peak stress, particularly in the longitudinal direction. Moreover, Figures 8 F and H depict the extent of tissue recovery per cycle, normalized to the second cycle. Both native and decellularized tissues demonstrated a consistent decline in recovery rate as cycles progressed, aligning with expected viscoelastic deformation.

**Figure 8.**
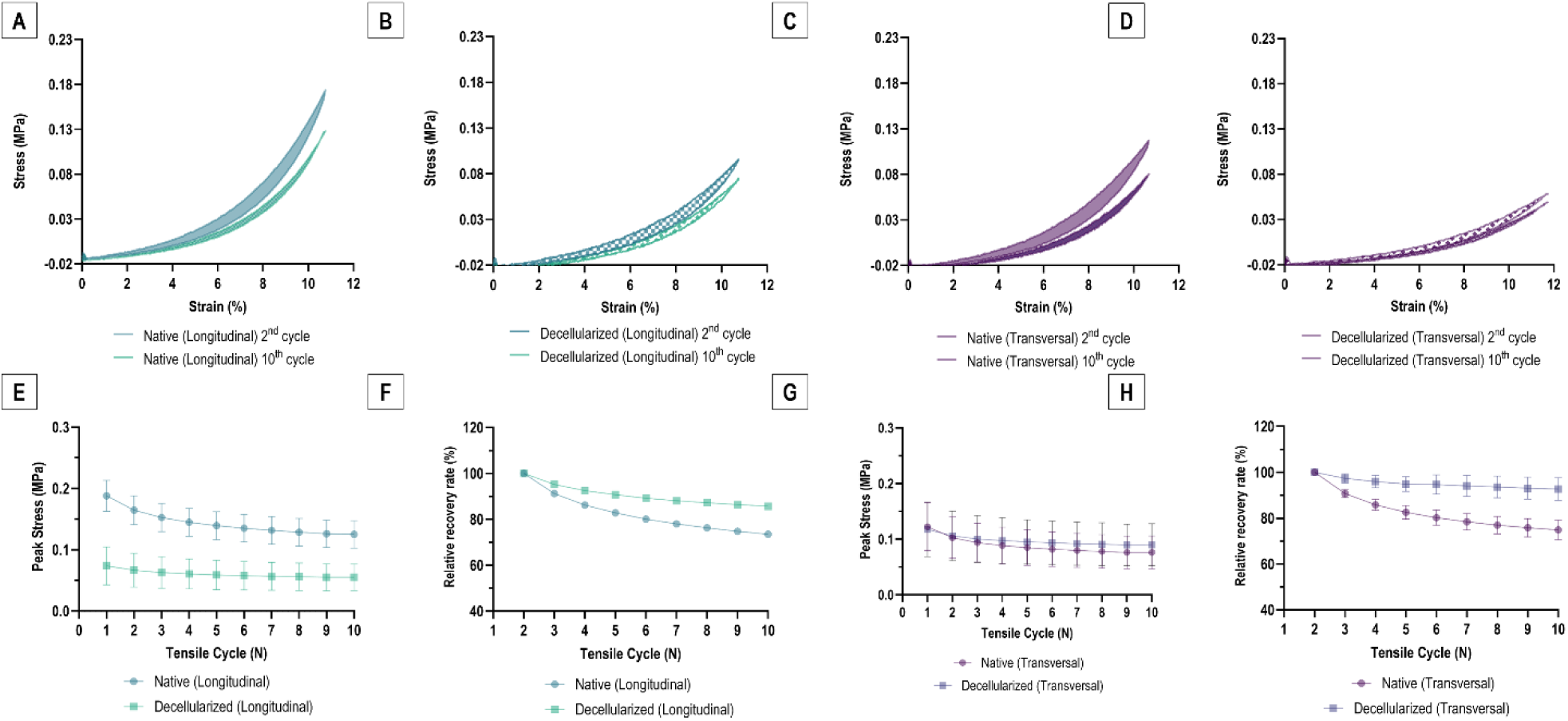
Cyclic mechanical behaviour of native and decellularized tissues. (A–D) Representative stress–strain curves obtained from cyclic uniaxial tensile testing for native and decellularized tissue in the longitudinal and transversal direction. (E, G) Peak stress response as a function of cycle at a constant applied strain. (F. H) Relative recovery rate per cycle in relation to the second cycle. Data are presented as mean ± standard deviation (n = 4).

In all groups, the area of the loading–unloading hysteresis loop decreased as cycling progressed. This trend appears to reflect reduced energy dissipation as the tissue became mechanically preconditioned under repeated loading. Native tissues generally exhibited larger hysteresis areas than decellularized tissues, an effect that was most evident in the longitudinal direction and may point to higher intrinsic damping and viscous losses in the native matrix. In longitudinal samples, the normalized recovery rate in native tissue declined to approximately 73% by cycle 10. Decellularized samples, by contrast, showed a smaller reduction and remained at around 85–86%. In the transverse direction, the hysteresis area also decreased with cycle number in both groups, although the magnitude of change differed. Native transverse samples showed a clear decline to roughly 68–74% by cycle 10. Decellularized transverse samples changed little over the same range, remaining close to their initial values (approximately 92–98% at cycle 10). Together, these results suggest that native transverse tissue undergoes pronounced stress softening and a progressive reduction in dissipative capacity over the first cycles, whereas decellularized tissue maintains a largely cycle-independent hysteretic response. Overall, starker recovery rates are noted in the native tissue when compared with the decellularized tissue and greater differences in peak stress in the longitudinal direction were observed.

### 3.5. Barrier ability and degradation profile of tissues

To evaluate if the applied treatment hinders the ability of the tissue to perform a barrier against microbial invasion a microbial penetration assay was performed over 14 days (Figure 9 A to C). The results revealed that along a period of 14 days the native and treated tissue was able to prevent contamination of the underlying media, exhibiting a level of efficacy that was analogous to that of commercial band-aids. This demonstrates that the decellularization protocol did not disrupt the fibres to the extent of allowing microbial invasion.

**Figure 9.**
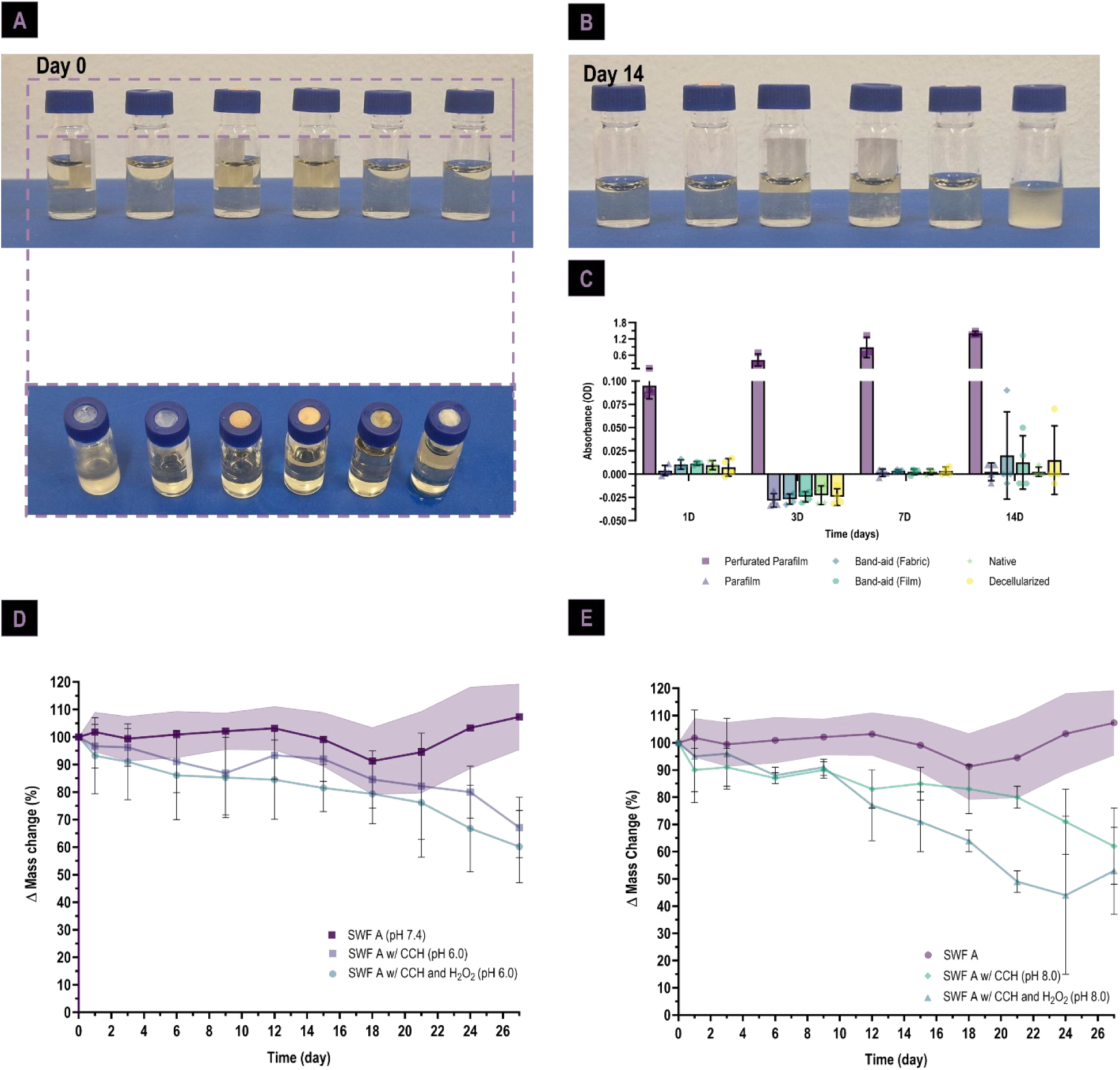
Biological assays on the native and treated tissue: microbial penetration assay at (**A**) 0 day with a top view of assay set up, (**B**) 14 days and (**C**) turbidimetry of microbial penetration over 14 days. Degradation assay: under (D) 6.0 and (E) 8.0 pH conditions with baseline degradation using SWF A at pH 7.4

Another important property of such an ECM-based patch is its ability to function as temporary or semi-permanent scaffold guiding wound regeneration, with a degradation profile designed to facilitate cell attachment, tissue remodelling and eventually total integration. The experimental setup encompassed four distinct scenarios, each involving a different acute environment (Figure 9 D to E). The results of the study indicate that SWF A baseline does not promote tissue degradation. Furthermore, the incorporation of collagenase at a pH of 6.0 results in a 33% mass reduction in a less challenging environment with respect to enzymes, while a 7% mass decrease is observed with the addition of oxidative stress over a period of 27 days (Figure 9 D). Moreover, an increase in the enzymatic challenge resulted in a 39% mass loss, with an additional 8% decrease occurring due to the addition of oxidative stress over a period of 27 days (Figure 9 E).

### 3.6. Surface screening through cell interaction

The decellularised small intestine exhibits two exposed surfaces with distinct topographies. Consequently, a screening was performed to ascertain which surface could have better potential for cell adhesion and proliferation using HDF and HACATs (Figure 10). As demonstrated in Figure 10A and B, human dermal fibroblasts exhibited a consistent increase in metabolic activity over a period of 14 days in both the submucosa and serosa surfaces. However, the proliferation rate was found to be generally higher in the serosa layer, particularly on the 14th day. As demonstrated in Figure 10 C and 10 D, human immortalised keratinocytes exhibited elevated metabolic activity in the serosa layer. The proliferation rates observed in both membranes were comparable.

**Figure 10.**
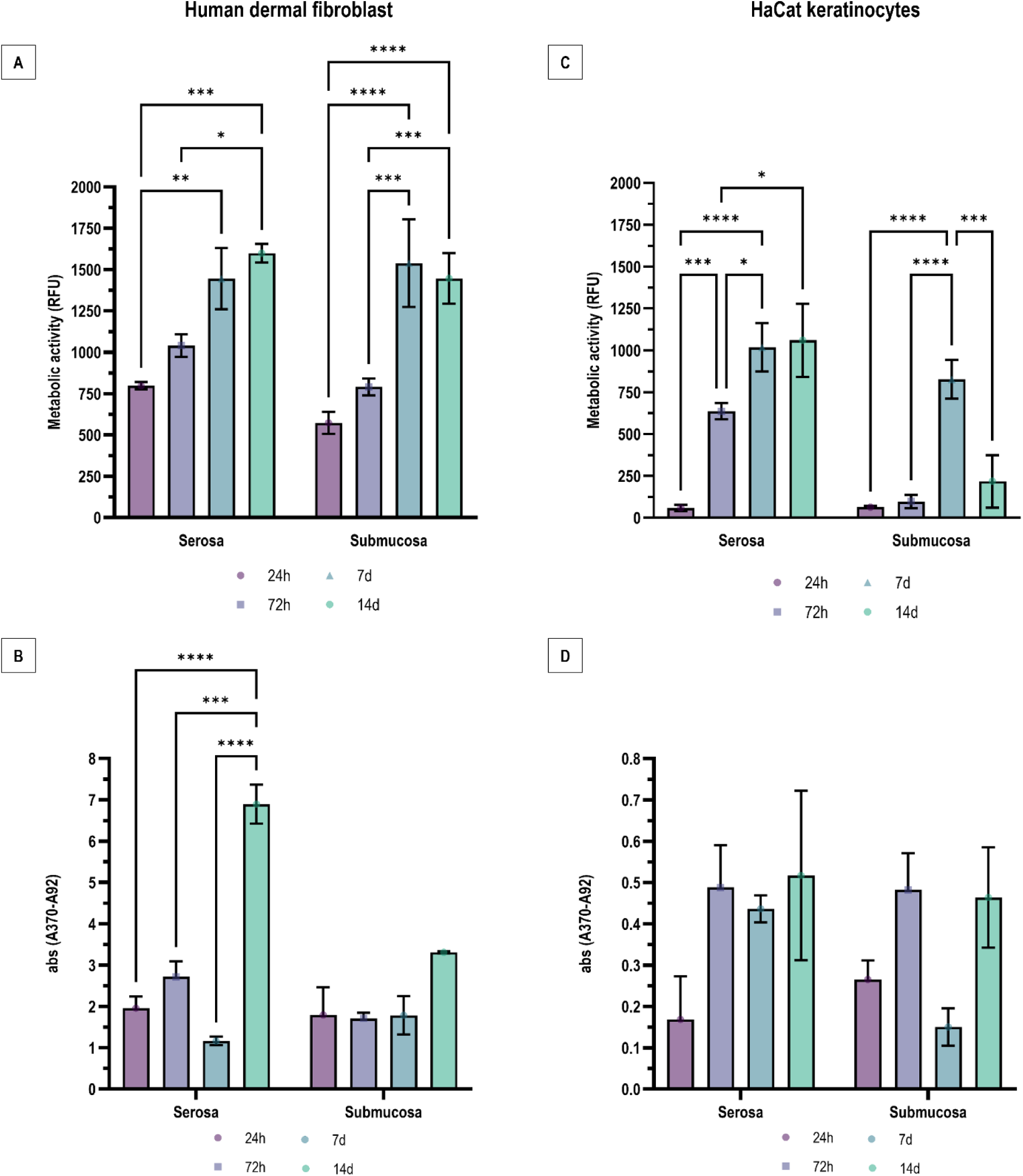
Screening of cell interaction with tissue surface of HDF (A,B) and HACAT (C,D) by metabolic activity and proliferation quantification.

## 4. Discussion

In this work, we looked at using DMSO as a penetration enhancer to decellularize full-thickness porcine small intestine and considered its possible applications for burn wounds. By using the entire intestinal wall rather than an isolated layer, the tissue seems to retain stronger mechanical properties and preserve much of its natural barrier function, features that could be particularly important for protecting a wound and supporting the healing process. Working with full-thickness ECM also provides a chance to compare the submucosa and serosa surfaces, which may differ in how suitable they are for direct wound contact. Interestingly, our approach removed cellular material effectively while largely keeping the extracellular matrix intact, along with the tissue’s structure, biochemical makeup, and functional characteristics. Taken together, these observations hint that full-thickness intestinal ECM might offer some advantages over thinner or partial segments when designing scaffolds for wound repair, though further studies will be needed to explore this fully.

Macroscopic observation revealed no obvious changes in tissue size or shape before and after treatment. Interestingly, the tissue presents a degree of transparency, likely due to the removal of blood and the thinness of the intestinal wall, which allows visualization through the membrane. This property may be advantageous for wound patches, as it enables real-time monitoring of the injured site without compromising coverage. Histological analysis confirmed efficient nuclear clearance, with residual DNA levels below the commonly accepted threshold of 50 ng mg⁻¹ ECM (10). The extensive nuclei removal observed in H&E staining suggests that DMSO can disrupt cell membranes effectively in combination with detergents, while also helping to reduce the immersion time in harsh detergents to about 9 hours, shorter than typical protocols (above 32h) that are applied to decellularize SIS membrane which is thinner that the whole SI (34–36). A secondary enzymatic step, such as DNase treatment, could potentially further shorten washing times and reduce PBS and ddH_2_O use, offering both efficiency and a more friendly-environmental approach. Notably, the ECM architecture and composition were preserved better than in some traditional methods (37, 38). SEM images showed intact villi and abundant coiled collagen fibers in the serosa, whereas histology indicated that overall fiber organization was retained in the serosa but particularly disrupted in the muscularis layer. This disorganization likely reflects the harsh nature of SDS/SDC, which are known to solubilize and damage matrix components, especially elastin and glycosaminoglycans (GAGs) (25, 26). Biochemical assays supported these observations: decellularized tissue retained approximately 44% of native GAGs but showed a significant reduction in alpha-elastin levels (42%). This pattern aligns with previous reports that demonstrate that elastin is more susceptible to degradation than collagen and is often preferentially lost during decellularization (39, 40). Similarly, tissues with elastin defects in pathological contexts tend to show accelerated enzymatic breakdown and fiber fragmentation relative to collagen (41). Amino acid profile is consistent with exposure of ECM collagen and other matrix proteins once cells and soluble components are removal. For example, Cys, a key crosslinkable residue, and abundant collagen-associated residues such as Gly, Ala, and Pro often dominate decellularized ECM profiles (42). Moreover, the profile is also consistent with preferential loss of intracellular proteins and more labile amino acids (43). In essence, the decellularization effectively removes cellular components while concentrating ECM-rich, collagen-associated amino acids, likely because the blood and cell residues contribute substantially with weight without adding significant content.

The WVTR remained comparable to human skin (≈200–300 g/m²·day), indicating that intrinsic matrix permeability is largely preserved, which similar to skin can allow sufficient vapor escape to prevent maceration while maintaining a balanced moist wound environment, essential in supporting re-epithelialization and healing (44, 45). These values fall within reported ranges for hydrogel and polysaccharide dressings used in burn and highly exuding wounds (46, 47). Water sweeling in decellularized matrices is largely governed by negatively charged GAGs and proteoglycans. Here, the decellularized intestine maintained effective moisture management while interacting with protein-rich wound exudates. Swelling studies seem to reflect a more preserved matrix property. In PBS, both native and decellularized tissues exhibited rapid initial swelling followed by a plateau, with similar equilibrium values, suggesting that overall hydrophilicity and pore architecture remained largely intact. In simulated wound fluid (SWF), swelling stabilized more slowly and reached higher equilibrium in decellularized tissue. This behaviour likely results from progressive protein adsorption mechanisms, which modifies surface chemistry and enhances fluid retention through interactions with collagenous scaffolds (19, 46, 48, 49). The observed difference underscores the increased accessibility of binding sites and internal surface area following cellular removal, suggesting the scaffold can absorb proteinaceous wound exudate effectively while maintaining WVTR conducive to moist wound healing (19, 46, 48, 49).

Mechanical testing reflected compositional and microstructural changes. Decellularized small intestine preserved inherent anisotropy, with significant greater stiffness along the longitudinal axis compared with the transverse direction, consistent with predominant collagen fiber alignment along the intestine reported for native and SIS tissues and seen in this work in histological sections (50–52). In contrast, transverse loading allowed more deformation at lower stress, likely due to fiber rotation and uncrimping rather than direct fiber stretching, mirroring previously reported nonlinear, anisotropic behaviour of intestinal wall and SIS scaffolds (51–54). Overall, longitudinal stiffness increased despite elastin loss, possibly due to enhanced crosslinking of the collagen/elastin architecture that governs load transfer induced by detergents and peracetic acid (51–53, 55). Cyclic loading revealed pronounced hysteresis and a gradual decline in peak stress, characteristic of preconditioning and viscoelastic softening seen in soft tissues such as skin and vessel walls (56–60). Notably, decellularized tissue exhibited more stable cyclic behaviour than native samples, which is consistent with reports that cellular and soluble components contribute substantially to viscoelasticity and inelastic deformation in soft tissues, and their removal yields a collagen-dominated matrix with reduced stress-softening and more predictable behaviour under repeated loading (26, 56, 61, 62). This stabilization is likely advantageous for long-term load-bearing applications, such as deep wound coverage with possible tissue integration.

Barrier function is another critical property for wound patches. Microbial penetration assays showed that both native and decellularized tissues prevented contamination of the underlying over 14 days, comparable to a commercial band-aid. Preservation of ECM continuity and density appears sufficient to block microbial invasion, a particularly important feature for burn and open wounds, where loss of the native skin barrier significantly increases infection risk (63).

Degradation studies further support the scaffold’s suitability for full-thickness burns. Healing of full-thickness burns is characterized by an initial phase dominated by tissue necrosis and inflammation, during which enzymatic activity at the wound surface is relatively limited, followed by a delayed remodelling phase associated with increasing protease activity and granulation tissue formation. In this context, wound patches are required to maintain structural integrity during early stages of healing while becoming progressively permissive to degradation and remodelling at later time points (64, 65). Minimal mass loss in SWF suggests stability during the early inflammatory phase with limit enzymatic challenge, while collagenase exposure induced gradual degradation compatible with later remodelling. Moreover, oxidative stress had only small effect, suggesting resistance to premature degradation in inflammatory conditions. The biphasic pattern, initial loss of soluble or weakly bound components followed by slower degradation of robust fibrillar ECM, aligns with time-dependent matrix remodelling rather than uncontrolled breakdown. Removal of cellular material increases internal surface area and pore accessibility, facilitating early enzyme interaction with exposed collagen, proteoglycan domains and easier to degrade ECM components. This initial phase is followed by a plateau, indicating that the remaining scaffold consists predominantly of more densely packed, fibrillar, and structurally robust ECM components that are less susceptible to rapid enzymatic cleavage.

The decellularized intestine presents two very structurally distinct surfaces revealed by SEM imaging. A collagen-rich serosa and a relatively smooth submucosa despite villi-related surface area increases. Historically, SIS has been favoured over the whole small intestine as a biomaterial, largely because isolating the submucosal layer produces a thin, ECM-rich scaffold with relatively low immunogenicity that is easier to standardize and adapt across different applications (11, 66–69). By contrast, retaining the full intestinal wall brings along mucosal, muscular, and neural components, which substantially increase cellular content, immunological risk, and structural heterogeneity. That added complexity does not just complicate decellularization; it also raises practical hurdles for quality control and regulatory approval. From a processing standpoint, SIS can be decellularized in a reproducible way using a range of physical–chemical protocols that reliably meet commonly accepted DNA thresholds (70). Even so, recent reviews point to considerable variability in SIS preparation and emphasize the need for better standardization and reporting (69). Extending that level of control to the full small intestine wall would likely be even more challenging. Part of SIS’s appeal is also mechanical and practical. Its natural thinness, flexibility, and suturability make it easy to handle, whether used as a single sheet, stacked into multilayer membranes for skin or abdominal wall repair, or rolled into conduits for applications such as urinary diversion (66, 67, 71). Over time, these properties—combined with extensive preclinical and clinical use in wound matrices, hernia repair, and urogenital or cardiovascular patches—have established a strong translational and regulatory track record for SIS that whole-SI constructs simply do not yet have lack (71). Standard SIS processing discards the muscular and vascular components of the intestine, even though these elements could be relevant for addressing one of the persistent challenges in tissue engineering: vascularization of larger constructs (72).

Using the full SI wall instead of just the submucosa could also, at least in principle, offer additional collagen-rich layers and greater overall thickness, which may be beneficial for mechanically demanding applications such as skin repair. Collagen is the primary load-bearing component of the dermis, and its dense, fibrillar organization underpins the tensile strength and resistance to deformation seen in native skin (73). Many collagen-rich tissues are, in fact, layered composites, with different strata contributing distinct mechanical roles (74). While the submucosa is already collagen rich, the intestinal wall as a whole contains additional fibrous collagen within the muscular and serosal regions, which help the native organ withstand internal pressure and tensile forces (74, 75). Preserving more of these layers would be expected to increase stiffness and thickness relative to SIS alone, much like the way full-thickness skin matrices outperform isolated collagen films in mechanical tests (74). For skin repair in particular, this higher-order collagen architecture may better approximate the mechanical “envelope” of native tissue than thin SIS membranes, which often need to be multilayered, chemically crosslinked, or combined with reinforcing materials to achieve sufficient strength (67). In practice, many SIS-based wound dressings already rely on added support—such as polymer nanofibers or inorganic fillers—to compensate for limited toughness in full-thickness defects (67, 76), suggesting that alternative strategies, including fuller use of the intestinal wall, may be worth revisiting.

In this work we found that human dermal fibroblasts proliferated preferentially on the serosa, likely due to dense collagen networks which provides ample adhesion points, whereas keratinocyte proliferation was similar on both sides, that could indicate re-epithelization. This indicates surface-specific bioactivity, which can provide optimal performance when using the serosa surface and when applied in a certain scaffold orientation.

In this work, a tissue-specific protocol has been developed to produce scalable decellularized intestinal patches that can preserve ECM structure, mechanical behaviour, and cellular compatibility, key factors for functional integration in skin grafts (77). Yet, xenogeneic considerations remain; for example, porcine α-Gal antigens must be evaluated to minimize immune rejection. One possible direction for further developing this dECM might involve incorporating antibacterial agents or particles, ranging from conventional antibiotics to metal or metal oxide nanoparticles, chitosan, copper ions, microbiota-derived postbiotics, or even nanozymes, by taking advantage of the matrix’s natural porosity, hydrophilicity, and available binding sites. This approach could likely allow both rapid and more sustained local release of these agents (78–84). By embedding antimicrobials directly within the dECM, the scaffold could actively manage infection at the wound surface, potentially lowering bacterial load and slowing biofilm formation without needing extra medicated dressings or heavy reliance on systemic antibiotics. Of course, whether this level of control is consistently achievable in complex, real-world wounds remain uncertain. Adjusting the degradation rate of the dECM and the interactions between drug and matrix may further allow fine-tuning of release profiles, high initial concentrations could knock down the bioburden quickly, followed by a longer, lower-level release that might help prevent reinfection throughout the healing process (85). Future work should also explore *in vivo* performance in burn models to confirm long-term efficacy.

## 5. Conclusion

In this work, we evaluated whether full-thickness porcine small intestine could be effectively decellularized using a DMSO-assisted protocol and assessed its suitability for wound healing applications. Working with the intact intestinal wall added a layer of difficulty, literally and structurally, yet the approach proposed achieved efficient cellular removal, with reduced residual DNA and largely preserved extracellular matrix architecture. Structural, biochemical, and mechanical analyses showed that the dSI remained predominantly collagen-based, anisotropic, and mechanically stable, although partial losses of elastin and glycosaminoglycans were observed, consistent with the use of ionic detergents. Several properties relevant to skin and burn wound repair were retained after decellularization. Water vapor transmission rates fell within ranges reported for human skin and clinical wound dressings, indicating the ability to maintain a moist wound environment without excessive occlusion. The matrix also resisted microbial penetration and showed limited mass loss under conditions mimicking the early wound phase, while remaining susceptible to enzymatic degradation associated with later remodelling stages. Taken together, this pattern points to a scaffold that is stable when protection is most critical, yet degradable accompanying cell remodelling. A notable finding was the functional difference between the two exposed surfaces of the dSI. The collagen-dense serosa side supported higher dermal fibroblast proliferation, whereas keratinocyte growth was comparable on both surfaces, which suggests that scaffold orientation may influence cell response and could be leveraged as a design parameter in wound applications. Overall, these results indicate that retaining the full intestinal wall, rather than isolating SIS, may offer advantages in terms of thickness, mechanical behaviour, and functional versatility, although *in vivo* studies will be required to assess translational relevance.

## Acknowledgements

This research was supported by National Funds from Fundação para a Ciência e a Tecnologia (FCT), through the Doctoral Research Grant 2021.05919.BD, Be@t-Textile Bioeconomy (TC-C12-i01, Sustainable Bioeconomy No. 02/C12-i01.01/2022) project, promoted by the Recovery and Resilience Program (RRP), Next Generation EU, 2021-2026, and the project IBEROS+ (0072_IBEROS_MAIS_1_E,Interreg-POCTEP 2021-2027), the International Cooperation Fund of the Science and Technology Commission of Shanghai Municipality (22520711900) are also acknowledged. The authors would like to thank Seara S.A. for providing porcine small intestine tissue used in the decellularization process.

